# The Standard Genetic Code Lacks Redundancy For Amino Acid Codons

**DOI:** 10.1101/2022.12.27.519817

**Authors:** Alan M. Laibelman

## Abstract

The Standard Genetic Code contains sixty-one codons used to translate twenty amino acids. A 3:1 ratio is presumed to mean all of them except methionine and tryptophan exhibit redundancy (also known as degeneracy) such that multiple codons are interchangeable with respect to cognate amino acids. Since the process of translation converting codon triplets into amino acids involves a 1:1 relationship between mRNA-containing codons and tRNA-containing anticodons, any physicochemical variability in tRNA molecules associated with the same amino acid must break redundancy among codons. By studying the characteristics of 186 tRNA sequences from Archaea, it is demonstrated that multiple features lead to the conclusion that every anticodon triplet is unique, and therefore that every codon triplet must likewise be unlike all others, even if coding for the same amino acid. Many implications necessarily accrue from this result related to the mechanism for translation, codon usage, and the evolution of the genetic code, all of which are discussed.

## INTRODUCTION

If any aspect of genetics has attained universal agreement, it is that based on translation of groups of three nucleotides (a codon triplet) from long mRNA strings, a standard code comprised of a set of sixty-four triplets, including three Stop codons, leads to a collection of twenty amino acids needed to satisfy the physical and biochemical needs of living things (Koonin and Novozhilov 2009). Long after this doctrine was enshrined as the *ne plus ultra* of genetics, relatively minor variations among Archaea, Bacteria, Eukarya, as well as some found in organellar genomes such as mitochondria and chloroplasts were discovered. Although this cumulative set of deviations has meant construction of several translation tables from the original, departures from the initial scheme did not introduce any profound modifications to the original thesis. The existence of a singular edifice remains unquestioned biological dogma even after it was revealed racemase enzymes enable formation of (D) amino acids from (L) stereoisomers and are crucial for biological function (Grishin *et al*. 2020), or that the existence of specialized atypical amino acids selenocysteine and pyrrolysine required expansion of the original twenty towards acceptance of at least twenty-two (Lin *et al*. 2017).

The genetic code assigns eighteen of the foundational twenty amino acids to remarkable displays of redundancy: twofold degenerate asparagine, aspartic acid, cysteine, glutamic acid, glutamine, histidine, lysine, phenylalanine, tyrosine; threefold isoleucine; fourfold alanine, glycine, proline, threonine, valine; sixfold arginine, leucine, serine. *This paper rejects redundancy as a logically implausible thesis from an evolutionary perspective, and untrue as a practical result of mechanical processes subsumed under the heading of translation*. It is accepted that the code employs multiple mRNA codons stipulating usage of the same amino acid during protein synthesis. Rejection of redundancy is based on denying that the tRNA molecules responsible for preparing amino acids for introduction into proteins are identical. Transfer RNA anticodon triplets forming base pairs with mRNA codon triplets are not isoacceptors for any amino acid. As a shorthand descriptor for an entire tRNA molecule, all anticodon triplets, original or later reassigned, are unique and distinguishable from each other.

Transfer RNA molecules are crucial to a translation process designed to construct multitudes of diverse proteins with a wide range of *in vivo* function. Does it make sense for every organism to devote precious, and quantitatively limited, energetic and/or material resources in fulfillment of a commitment to produce multiple forms of the same tRNA? Multiple *forms* are not equivalent to multiple *copies*. The first refers to anticodons functionally indistinguishable *in vivo*; the second designates tRNA duplicated exactly to guard against damage or mutation inducing permanent disability (Berg and Brandl 2021).

There are no duplicate biosynthetic pathways to produce the amino acids themselves (Nelson and Cox 2021, Ch 22). Why should there be redundant modes for their employment through the genetic code? Do duplicate Tricarboxylic Acid cycles (Nelson and Cox 2021, Ch 16) exist, much less sixfold degenerate sets of identical cycles? Is there a second electron transfer metabolic redox pathway (Nelson and Cox 2021, Ch 13), much less a sixfold degenerate group of chemical sequences serving the same purpose?

Having a backup plan is *a priori* rational in case of breakdown; for example, there is inherent capability for reassignment of nerve function after disease or injury. However, neuroplasticity involves switching across initial purposes, not an originally designed clone available from the lifeform’s inception (Voss *et al*. 2017).

From one perspective, evolution is reproducibly constant over long stretches of cosmic time because biochemical procedures controlled in large part by the genetic code are nearly invariant. This internal consistency led to the notion of a *minimal gene set* applicable to all creatures, yet still permitting classification as Archaea, Bacteria, Eukarya (Koonin 2000; Charlebois and Doolittle 2004). From another point of view, evolution must permit flexibility so there is freedom to adapt to changing environmental conditions over that same period of cosmic time. Evolutionary adjustments imply recognizable alterations to biochemical pathways enabling initial development of organisms, so genetic codes must be constructed to embody access to required variation (Caetano-Anollés *et al*. 2013). Complementary constructs of code-inspired mechanistic constancy and permissive variability in end products undergird *phylogeny*, whose central thesis is the postulated existence of a Last Universal Common Ancestor unifying the three life domains (Glansdorff *et al*. 2008).

Access to flexibility argues against employing degeneracy as an optimal operational modality. To build only to break down and reorient, if necessary, is inefficient; organisms literally cannot usually afford to be so wasteful (LaRowe and Amend 2016). Catastrophe-induced neuroplasticity is proof that structural realignment is an emergency procedure, not par for the course. Genetic realignment would likewise be an emergency procedure *if* it had not already been ingrained in the genetic code. The major aim of this paper is to provide some details illuminating this innate ability to genetically realign as needed.

One pathway for biochemical versatility is through post-translational modification of amino acids (Bohlke and Budisa 2014). Thousands are described and made publicly available through dbPTM (http://awi.cuhk.edu.cn; Li *et al*. 2022). Similarly, nucleotides have undergone pre-translational changes in tRNA (Grosjean *et al*. 2010). These approaches to adaptability are productively beneficial, but speak against translational redundancy as compelling.

A second mechanism for addressing flexibility is via externally-sourced mutation of nucleotides. (Loewe and Hill 2010). Mutations, almost by definition, occur haphazardly by positioning randomly in genomic sequences over uncontrolled timespans, not as products of intentional introduction leading to a desired end. Consequently, most mutations are harmful and fairly quickly excised from gene pools. It is the case that a small number are benign, yet tolerated because energy to be expended in removing them and reverting to the original versions does not lead to acceptable cost/benefit estimations. This situation shows that organisms do not ordinarily seek to expend energy unless deemed crucial to survival. Lastly, there develops a relatively minuscule number of useful mutations retained and reproduced in subsequent generations. However, the rarity of its occurrence implies it is not an efficient, dependable, generalizable methodology for any species to invoke as an ordained pathway for flexibility.

A third natural route to genomic diversity is through horizontal gene transfer: steal what is needed from an organism that has already solved the problem(s) encountered. At a time when just seven archaeal genomes were completely sequenced, it was estimated that 5−14% of all genes were acquired through this process (Garcia-Vallvé *et al*. 2000). It is a highly effective strategy in many instances, known to transpire in, and across, all three domains (Keeling and Palmer 2008). However, it is uncertain whether adequately generalizable to all situations. From a research perspective, when encountering horizontal gene transfer events, it is often difficult to determine whether A borrowed from B, B borrowed from A, or both obtained their material independently from third lifeform C (Sevillya *et al*. 2020).

What is proposed elucidates a channel to genomic adaptability in use, but unrecognized. *Codon redundancy does not exist because there are no redundant base paired anticodons; each possesses unique characteristics important for translation*. A cultural analogy would be to specialized roles for athletes on baseball, football, or basketball teams. For termination codons UAA, UAG, UGA, individuality has been attributed: (i) ribosomal complexes interact with different release factors (Korkmaz *et al*. 2014); (ii) they vary in ability to be reassigned to novel amino acids, such as UGA for selenocysteine (Rother *et al*. 2000) or UAG for pyrrolysine (Brugère *et al*. 2018). Moreover, isoleucine’s AUA codon is usually (there are species exceptions) reassigned: the expected tRNA anticodon is not UAU, but C^+^AU (C^+^ = agmatidine) in Archaea (Mandal *et al*. 2010) or LAU (L = lysidine) in Bacteria (Köhrer *et al*. 2014).

These instances are *not* illustrative of the argument against codon redundancy being suggested. Consider: the Periodic Table of Elements contains far more element variation than noted in the basic chart because listed atomic masses are actually weighted averages of all isotopic masses represented by a single chemical symbol. Analogously, each anticodon triplet actually conveys a set of identities revealed in part through the number of nucleotides in tRNA molecules assigned to each amino acid, and in part through the sequence of an entire string. By analyzing variety in tRNA employed, a profile unique to each codon/ anticodon pair is obtained. Anticodon uniqueness necessarily implies codon novelty, so no redundancy in the applicable genetic code for each lifeform.

Variations in tRNA do not extend to aminoacyl-tRNA synthetases catalyzing formation of tRNA-to-amino acid chemical bonds because synthetases are usually found in each specimen as a single enzyme copy per amino acid (Chaliotis *et al*. 2017), although exceptions exist where 0−2 copies are incorporated. Lysine is unusual in that its synthetase comes in two versions differing in placement within the ribosome complex, and more specifically, in their interaction with tRNA (de Farias and Guimarães 2008).

## RESULTS

Comparative genomics studies endorse two assumptions: (i) analyses depend on accurate genome sequencing by researchers without introducing errors during result tabulation; (ii) collections of sequences stored in public databases are faithful expressions of original data without creating errors during receipt, storage, or release. Published sequence information is rarely independently replicated; hence, conclusions drawn by those other than the initial investigators rely upon that information’s *a priori* correctness for subsequent analysis. These epistemological conditions are upheld in the present case: all evaluations, as well as their significance, are grounded on implied correct sequencing information found in GtRNAdb (http://gtrnadb.ucsc.edu; release 19, June 2021; Chan and Lowe 2009).

Nucleotides modified pre-translation from the fundamental A/C/G/U options will not be factored into tRNA sequence variations except for anticodon C^+^AU symbolizing reassigned tRNA^2Ile^. Justification for exclusion of this complicating feature derives from a lack of knowledge concerning such alterations in the majority of Archaea (Wolff *et al*. 2020).

As a preliminary illustration of legitimacy in suggesting the genetic code is not redundant, despite all profession to the contrary, six Eukarya and six Bacteria were chosen quasi-randomly and evaluated for the number of nucleotides contained by every tRNA molecule transcribed by each organism. Results for the conventional sixty-one amino acid-specifying codons are produced in Table 1 for Eukarya and Table 2 for Bacteria. Species are arranged alphabetically by genus, and anticodon triplets ordered by three-letter amino acid designations. Transfer RNA lengths are measured in nt units. For twelve genomes, several points demand notice:

**Table 1.** Number of Nucleotides in tRNA molecules for Six Eukarya.

| AA | Anticodon | Alligator mississippiensis | Arabidopsis thaliana | Drosophila melanogaster | Gallus gallus | Homo sapiens | Saccharomyces cerevisiae |
| --- | --- | --- | --- | --- | --- | --- | --- |
| Ala | AGC | 73, 74 | 73 | 73 | 73 | 72, 73 | 73 |
|  | CGC | 72, 73 | 73 | 72 | 72 | 72 |  |
|  | UGC | 72, 73 | 73 | 72 | 72 | 71, 72 | 73 |
| Arg | ACG | 73 | 73 | 73 | 73 | 73 | 73 |
|  | CCG | 73 | 73 |  | 73 | 73 | 72 |
|  | UCG | 73 | 74 | 73 | 73 | 73 |  |
|  | UCU | 71, 72, 73, 74 | 73 | 73 | 73, 74 | 73, 74 | 72 |
|  | CCU | 73 | 73 | 73 | 73 | 73 | 72 |
| Asn | AUU | 73 |  |  |  |  |  |
|  | GUU | 74, 82 | 72, 74, 75 | 74 | 74 | 74, 75 | 74 |
| Asp | GUC | 71, 72 | 72 | 72 | 72 | 72 | 71, 72 |
| Cys | ACA | 72, 73 |  |  |  |  |  |
|  | GCA | 72, 73, 81 | 72 | 72 | 72 | 70, 72, 73 | 72 |
| Gln | CUG | 72 | 72 | 72 | 72 | 72 | 72 |
|  | UUG | 72, 73 | 72 | 72 | 72 | 72 | 72 |
| Glu | CUC | 71, 72 | 73 | 72 | 72 | 72 | 72 |
|  | UUC | 72 | 72 | 72 | 72 | 72 | 72 |
| Gly | CCC | 71 | 71 |  | 71 | 71 | 72 |
|  | GCC | 71 | 71 | 71 | 71 | 71 | 71 |
|  | UCC | 71, 72 | 72 | 72 | 72 | 72 | 72 |
| His | GUG | 72 | 72 | 72 | 72 | 72 | 72 |
| Ile | AAU | 74 | 74 | 74 | 74 | 74 | 74 |
|  | GAU |  |  |  |  | 74 |  |
|  | UAU | 74 | 74 | 74 | 74 | 74 | 73 |
| Leu | AAG | 72, 82 | 72, 81 | 82 | 82 | 82 |  |
|  | CAG | 72, 83 | 81 | 83 | 82, 83 | 83 |  |
|  | GAG |  |  |  |  |  | 82 |
|  | UAG | 82 | 80 | 80 | 82 | 82 | 82 |
|  | UAA | 83, 84 | 83, 84 | 84 | 83 | 83 | 84 |
|  | CAA | 84 | 84 | 83 | 83, 84 | 74, 83, 84 | 82 |
| Lys | CUU | 73 | 73 | 73 | 73 | 73, 74 | 73 |
|  | UUU | 73, 77 | 72 | 73 | 73 | 73 | 73 |
| Met | CAU | 73 | 74 | 73 | 73 | 73 | 73 |
| iMet | CAU | 72 | 72 | 72 | 72 | 72 | 72 |
| Phe | GAA | 73 | 73 | 73 | 73 | 73, 74 | 73 |
| Pro | AGG | 72 | 72 | 72 | 72 | 72 | 72 |
|  | CGG | 72 | 72 | 72 | 72 | 72 |  |
|  | UGG | 72 | 72 | 72 | 72 | 72 | 72 |
| SeC | GCA | 87 |  |  |  |  |  |
|  | UCA | 87 |  | 87 | 87 | 87 |  |
| Ser | AGA | 82 | 82, 84 | 82 | 82 | 82 | 82, 83 |
|  | CGA | 82 | 82 | 82 | 82 | 82 | 82 |
|  | GGA |  | 87 |  |  |  |  |
|  | UGA | 82, 83 | 82 | 82 | 82 | 82 | 82 |
|  | GCU | 71, 72, 82 | 82 | 82 | 82 | 82, 84 | 81, 82 |
| sup | UCA | 73 |  |  |  |  |  |
| Thr | AGU | 73, 74 | 74 | 74 | 73, 74 | 74 | 73 |
|  | CGU | 72, 73, 74 | 72 | 72 | 72 | 72, 74 | 72 |
|  | UGU | 72, 73, 74 | 73 | 72 | 72, 73, 74 | 72, 73, 74 | 72 |
| Trp | CCA | 72 | 72, 74 | 72 | 72 | 72 | 72 |
| Tyr | AUA |  |  |  |  | 73 |  |
|  | GUA | 72, 73 | 73 | 73 | 73 | 72, 73 | 75 |
| Val | AAC | 73 | 74 | 73 | 73 | 72, 73 | 74 |
|  | CAC | 73 | 74 | 73 | 73 | 73 | 73 |
|  | GAC | 73 |  |  | 73 |  |  |
|  | UAC | 73 | 73 | 73 | 73 | 73 | 74 |

**Table 2.**
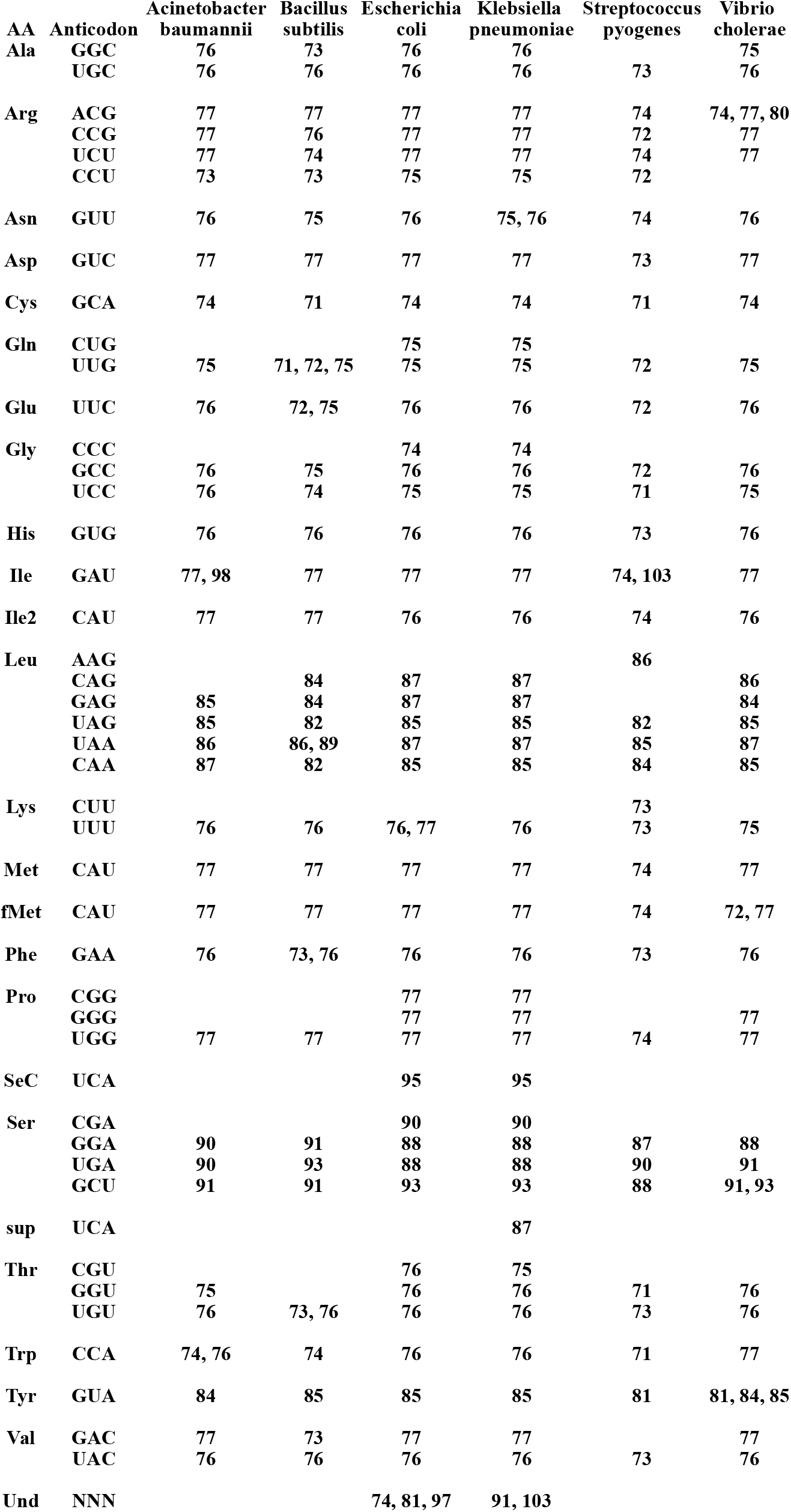
Number of Nucleotides in tRNA molecules for Six Bacteria.

| AA | Anticodon | Acinetobacter<br>baumannii | Bacillus<br>subtilis | Escherichia<br>coli | Klebsiella<br>pneumoniae | Streptococcus<br>pyogenes | Vibrio<br>cholerae |
| --- | --- | --- | --- | --- | --- | --- | --- |
| Ala | GGC | 76 | 73 | 76 | 76 |  | 75 |
|  | UGC | 76 | 76 | 76 | 76 | 73 | 76 |
| Arg | ACG | 77 | 77 | 77 | 77 | 74 | 74, 77, 80 |
|  | CCG | 77 | 76 | 77 | 77 | 72 | 77 |
|  | UCU | 77 | 74 | 77 | 77 | 74 | 77 |
|  | CCU | 73 | 73 | 75 | 75 | 72 |  |
| Asn | GUU | 76 | 75 | 76 | 75, 76 | 74 | 76 |
| Asp | GUC | 77 | 77 | 77 | 77 | 73 | 77 |
| Cys | GCA | 74 | 71 | 74 | 74 | 71 | 74 |
| Gln | CUG |  |  | 75 | 75 |  |  |
|  | UUG | 75 | 71, 72, 75 | 75 | 75 | 72 | 75 |
| Glu | UUC | 76 | 72, 75 | 76 | 76 | 72 | 76 |
| Gly | CCC |  |  | 74 | 74 |  |  |
|  | GCC | 76 | 75 | 76 | 76 | 72 | 76 |
|  | UCC | 76 | 74 | 75 | 75 | 71 | 75 |
| His | GUG | 76 | 76 | 76 | 76 | 73 | 76 |
| Ile | GAU | 77, 98 | 77 | 77 | 77 | 74, 103 | 77 |
| Ile2 | CAU | 77 | 77 | 76 | 76 | 74 | 76 |
| Leu | AAG |  |  |  |  | 86 |  |
|  | CAG |  | 84 | 87 | 87 |  | 86 |
|  | GAG | 85 | 84 | 87 | 87 |  | 84 |
|  | UAG | 85 | 82 | 85 | 85 | 82 | 85 |
|  | UAA | 86 | 86, 89 | 87 | 87 | 85 | 87 |
|  | CAA | 87 | 82 | 85 | 85 | 84 | 85 |
| Lys | CUU |  |  |  |  | 73 |  |
|  | UUU | 76 | 76 | 76, 77 | 76 | 73 | 75 |
| Met | CAU | 77 | 77 | 77 | 77 | 74 | 77 |
| fMet | CAU | 77 | 77 | 77 | 77 | 74 | 72, 77 |
| Phe | GAA | 76 | 73, 76 | 76 | 76 | 73 | 76 |
| Pro | CGG |  |  | 77 | 77 |  |  |
|  | GGG |  |  | 77 | 77 |  | 77 |
|  | UGG | 77 | 77 | 77 | 77 | 74 | 77 |
| SeC | UCA |  |  | 95 | 95 |  |  |
| Ser | CGA |  |  | 90 | 90 |  |  |
|  | GGA | 90 | 91 | 88 | 88 | 87 | 88 |
|  | UGA | 90 | 93 | 88 | 88 | 90 | 91 |
|  | GCU | 91 | 91 | 93 | 93 | 88 | 91, 93 |
| sup | UCA |  |  |  | 87 |  |  |
| Thr | CGU |  |  | 76 | 75 |  |  |
|  | GGU | 75 |  | 76 | 76 | 71 | 76 |
|  | UGU | 76 | 73, 76 | 76 | 76 | 73 | 76 |
| Trp | CCA | 74, 76 | 74 | 76 | 76 | 71 | 77 |
| Tyr | GUA | 84 | 85 | 85 | 85 | 81 | 81, 84, 85 |
| Val | GAC | 77 | 73 | 77 | 77 |  | 77 |
|  | UAC | 76 | 76 | 76 | 76 | 73 | 76 |
| Und | NNN |  |  | 74, 81, 97 | 91, 103 |  |  |

- No species individually utilizes all sixty-one possible anticodon triplets, a fact probably attributable to the wobble hypothesis (Crick 1966).
- The twelve species collectively do not incorporate all possible anticodons. This is not a function of the wobble hypothesis, but of the paucity of examples. Transfer RNA for Arg^(GCG)^, Asp^(AUC)^, Gly^(ACC)^, His^(AUG)^, Phe^(AAA^), Ser^(ACU)^ are missing. When Archaeal tRNA molecules are later included, omissions reduce to Asp^(AUC)^, His^(AUG)^, Phe^(AAA)^, Ser^(ACU)^. Absent anticodons might be recovered when extensive explorations of sequenced bacterial and/or eukaryl tRNA molecules are undertaken, since GtRNAdb includes sequences for 4037 Bacteria and 576 Eukarya. If all sixty-one are eventually found in use, then recourse to the wobble hypothesis as applicable rule to explain absences within any organism becomes less reasonable, even if it should still prove to be generally of operational validity.
- Most anticodons are encoded with tRNA of one length per genome; in a few cases, 2−3 lengths are involved; *Alligator mississippiensis* invokes four tRNA^Arg(UCU)^ (71−74 nt). If it is hypothesized such variance is functionally irrelevant, then it is incumbent upon purveyors of this proposition to provide justification rather than merely assert it as controlling doctrine.
- Additional explanations are needed to rationalize why nucleotide number variance for an anticodon triplet occurs forty-seven times in six Eukarya plus sixteen times in six Bacteria. The genetic code is believed to be evolved to a level of high fidelity for translation (Freeland *et al*. 2003). Variable tRNA length without real world meaning confers contrary notions: although the code is relatively pristine in function, it tolerates sloppiness in execution.

If the last argument chain remains unconvincing, notice that, within an organism, *tRNA charging the same amino acid display different lengths*. Tables 1−2 allow ready confirmation and permit observing how frequently this eventuates. One exemplar from each species suffices; numbers in parentheses reflect nucleotides in each tRNA string:

*Alligator mississippiensis* → Asn^(AUU)^ (73); Asn^(GUU)^ (74, 82)

*Arabidopsis thaliana* → Lys^(CUU)^ (73); Lys^(UUU)^ (72)

*Drosophila melanogaster* → Gly^(GCC)^ (71); Gly^(UCC)^ (72)

*Gallus gallus* → Leu^(AAG)^ (82); Leu^(CAG)^ (82, 83); Leu^(UAG)^ (82); Leu^(UAA)^ (83); Leu^(CAA)^ (83, 84)

*Homo sapiens* → Ala^(AGC)^ (72, 73); Ala^(CGC)^ (72); Ala^(UGC)^ (71, 72)

*Saccharomyces cerevisiae* → Ile^(AAU)^ (74); Ile^(UAU)^ (73)

*Acinetobacter baumannii* → Val^(GAC)^ (77); Val^(UAC)^ (76)

*Bacillus subtilis* → Arg^(ACG)^ (77); Arg^(CCG)^ (76); Arg^(UCU)^ (74); Arg^(CCU)^ (73)

*Escherichia coli* → Gly^(CCC)^ (74); Gly^(GCC)^ (76); Gly^(UCC)^ (75)

*Klebsiella pneumoniae* → Ser^(CGA)^ (90); Ser^(GGA)^ (88); Ser^(UGA)^ (88); Ser^(GCU)^ (93)

*Streptococcus pyogenes* → Leu^(AAG)^ (86); Leu^(UAG)^ (82); Leu^(UAA)^ (85); Leu^(CAA)^ (84)

*Vibrio cholerae* → Ala^(GGC)^ (75); Ala^(UGC)^ (76)

Summary data from twelve species is precursor to reporting a comprehensive analysis performed on tRNA molecules from 186 Archaea. GtRNAdb offers sequences for 217 Archaea, but thirty-one were rejected because designated by genus + strain instead of a recognizable genus + species nomenclature. The retained dataset includes multiple strains from one species, especially orders Methanosarcinales and Sulfolobales. Multiple strains complicate results because they frequently possess identical length tRNA per anticodon, with replication distorting the distribution of lengths.

Sixty-two strains for sixteen species: *Archaeoglobus fulgidus* (2); *Desulfurococcus amylolyticus* (2); *Haloarcula hispanica* (2); *Haloferax mediterranei* (2); *Haloquadratum walsbyi* (2); *Metallosphaera sedula* (2); *Methanobacterium formicicum* (2); *Methanococcus maripaludis* (5); *Methanosarcina barkeri* (5); *Methanosarcina mazei* (7); *Methanosarcina siciliae* (3); *Methanosarcina thermophila* (2); *Pyrococcus furiosus* (2); *Saccharolobus islandicus* (15); *Saccharolobus solfataricus* (5); *Sulfolobus acidocaldarius* (4).

Supplemental Table 1 produces a roster of included Archaea by name according to the prokaryote Genome Taxonomy Database (http://gtdb.ecogenomic.org; version R207, June 2021). Some taxonomic names in GtRNAdb have been reassigned by GTdb and regarded as definitive (e.g., *Sulfolobus islandicus* is now *Saccharolobus islandicus*). Table S1 gives name and strain, altered if necessary, along with NCBI assembly numbers (https://www.ncbi.nlm.nih.gov) because the latter do not change even if nomenclature does.

Supplemental Table 2 gives nucleotide sequence information for every tRNA encoded in each genome for 186 Archaea. It constitutes the raw data from which all analyses are extracted. Multiple copies for individual tRNA anticodon triplets are presented on separate lines. When occurring, copies usually number two or three per anticodon, although those for cysteine go as high as eight.

Table 3 summarizes the distribution of tRNA lengths per anticodon for all Archaea. Nucleotide number was provided by GtRNAdb and used without alteration. Enormous variation in length is obvious, with ranges differing for each anticodon. Table 3 thus complements Tables 1−2 for Eukarya and Bacteria, respectively.

**Table 3.** Number of Nucleotides in tRNA Molecules for 186 Archaea.

| Anticodon | tRNA length (in # nt) | Anticodon | tRNA length (in # nt) |
| --- | --- | --- | --- |
| Ala (CGC) | 66,72,73,74,75,76,77,78 | Lys (CUU) | 73,74,75,76,77,78 |
| Ala (GGC) | 72,73,74,75,76,77,78 | Lys (UUU) | 73,74,75,76,77,78 |
| Ala (UGC) | 72,73,74,75,76,77,78 | Met (CAU) | 73,74,75,76,77,78,79,80,109 |
| Arg (ACG) | 73 | iMet (CAU) | 74,75,76,77,78,79 |
| Arg (CCG) | 72,73,74,75,76,77,78,79 | Phe (GAA) | 70,71,72,73,74,75,76,77 |
| Arg (CCU) | 73,74,75,77,78,79 | Pro (CGG) | 72,73,74,75,76,77,78,79 |
| Arg (GCG) | 72,73,74,75,76,77,78,79,85 | Pro (GGG) | 71,73,74,75,76,77,78,79 |
| Arg (UCG) | 72,73,74,75,76,77,78,79 | Pro (NGG) | 78 |
| Arg (UCU) | 72,74,75,76,77,78,83 | Pro (UGG) | 71,73,74,75,76,77,78,79,105 |
| Asn (GUU) | 70,71,72,73,74,75,76,77,78 | SeC (UCA) | 89,90,91,92 |
| Asp (GUC) | 68,72,73,75,76,77,78 | Ser (CGA) | 53,57,61,65,74,80,82,83,84,85,86,87,88,89,90,96 |
| Cys (GCA) | 71,72,73,74,75,76,77,78 | Ser (GCU) | 70,81,82,83,84,85,86,87,88,89,91 |
| Gln (CUG) | 72,73,74,75,76,77 | Ser (GGA) | 81,82,83,84,85,86,87,88,89,91,109 |
| Und (NUG) | 76 | Ser (UGA) | 81,82,83,84,85,86,87,88,89,90,91 |
| Gln (UUG) | 64,71,72,73,74,75,76,77,90 | sup (UCA) | 74 |
| Glu (CUC) | 73,75,76,77,78,79,191 | Thr (CGU) | 72,73,74,75,76,77,78 |
| Glu (UUC) | 73,75,76,78,79,90 | Thr (GGU) | 71,72,73,74,75,76,77,78,88,102 |
| Gly (ACC) | 82 | Thr (UGU) | 69,70,72,73,74,75,76,77,78 |
| Gly (CCC) | 71,72,73,74,75,76,77,78,79 | Trp (CCA) | 71,72,73,74,75,77,78,79,98 |
| Gly (GCC) | 71,72,73,74,75,76,77,78,79,84 | Tyr (GUA) | 71,72,73,74,75,76,77,78 |
| Gly (UCC) | 70,71,72,73,74,75,76,77,78,79,127 | Val (CAC) | 72,73,74,75,76,77,78 |
| His (GUG) | 71,72,73,74,75,76,77,78,80 | Val (GAC) | 65,72,73,74,75,76,77,78 |
| Ile (GAU) | 73,74,75,77,78 | Val (UAC) | 72,73,74,75,76,77,78,80,108 |
| Ile (UAU) | 74,75 | ??? (NNN) | 56,63,64,67,69,72,73,74,77,78,79,87,89,98,99,102,107,109,113,151,182,193 |
| Ile 2 (C<sup>+</sup>AU) | 72,73,74,75,76,77,78 |  |  |
| Leu (AAG) | 85 |  |  |
| Leu (CAA) | 55,68,70,71,76,81,82,83,84,85,86,87,88,96,97,101,128,218 |  |  |
| Leu (CAG) | 83,84,85,87,88,125,227 |  |  |
| Leu (GAG) | 82,83,84,85,87,88,90 |  |  |
| Leu (NAG) | 88 |  |  |
| Leu (UAA) | 77,83,84,85,86,87,88,90 |  |  |
| Leu (UAG) | 64,83,84,85,86,87,88,90,127 |  |  |

Supplemental Table 3 expresses the distribution of tRNA nucleotide number for every anticodon triplet, offering detail for that diversity. Since the number of organisms employing each anticodon triplet also varies, Supplemental Table 4 covers identical information to Table S3 in the form of percentage of total sample number per anticodon; it can be conceived as a normalized version of Table S3. Frequency % per anticodon = [(# occasions of length L) / (# tRNA per anticodon)] × 100%. General features of this tRNA catalog:

- Smallest number of nucleotides encoded in any tRNA molecule is 53
- Largest number of nucleotides encoded in any tRNA molecule is 227
- Total number of encoded tRNA for 186 Archaea is 8,869; average = 47.7
- Most frequent nucleotide length is 75, with frequency % = 1617 of 8869 = 18.2%
- *Thermococcus A litoralis* contains three incompletely characterized sequences: tRNA^Gln(NUG)^ (76 nt); tRNA^Leu(NAG)^ (88 nt); tRNA^Pro(NGG)^ (78 nt), All are maintained as stand-alone entries in the Tables.
- *Methanosarcina mazei* strain Tuc01 is unlike the other six strains with respect to the total number of tRNA encoded in its genome: five present 57 tRNA, one codes for 59 tRNA, but strain Tuc01 has just 43 tRNA according to the GtRNAdb source.

If the apparent discrepancy for *Methanosarcina mazei* strain Tuc01 is authentic, it seems strange it would represent the same species as the other six strains. If this distinction constitutes a data error, it is to be determined whether the research team did not detect and sequence all tRNA in the genome, or failed to catalog all molecules found and characterized. It is also conceivable fault lies in data transfer to public databases, or that sequence recipients encountered reception and/or storage mishaps. Assumptions innate to genomics research were articulated earlier for the purpose of evaluating the merits of all options.

The data in Tables 3, S3−S4 unequivocally demonstrate distribution profiles for encoded tRNA lengths in 186 Archaea are unique for each anticodon triplet. Given the innate bonding complementarity for codon/anticodon pairs, inclusive of wobble base relationships where existent, this observation implies every corresponding codon triplet in the genetic code exhibits zero redundancy regardless of amino acid translated.

Some tRNA exhibit a *relatively flat* profile in that several lengths are of nearly equal frequency: tRNA^Gly(CCC)^ has distribution percentages for its three most common encoded lengths of 25.6% (71 nt), 25.6% (76 nt), 19.4% (72 nt). Other tRNA exhibit a *dominant peak* profile with a large disparity in frequency by length: tRNA^Asp(GUC)^ has distribution percentages for its three most commonly encoded lengths of 54.8% (73 nt), 18.9% (75 nt), 16.6% (78 nt). Notice that the preferred lengths for tRNA^Gly(CCC)^ have none in common with preferred lengths for tRNA^Asp(GUC)^; this is an indicator of diversity expressed by archaeal tRNA, and hinted at by the examples found for Eukarya and Bacteria in Tables 1−2. Stronger evidence for anticodon nonredundancy is offered by tRNA^Ser^ (four encoded in Archaea). The three most frequent tRNA string lengths (nt in parentheses) for each of the four:

- 175 tRNA^Ser(CGA)^ = 27.4% (85), 24.0% (84), 9.7% (87)
- 191 tRNA^Ser(GGA)^ = 24.6% (84), 14.1% (83), 13.6% (85)
- 185 tRNA^Ser(UGA)^ = 30.8% (85), 19.5% (84), 15.1% (87)
- 190 tRNA^Ser(GCU)^ = 51.6% (85), 10.0% (84), 9.5% (88)

Although the most frequent lengths are nearly the same in all cases, distributions by percentage reveal different patterns. The first three tRNA are characterized as *relatively flat*, but tRNA^Ser(GCU)^ is of the *dominant peak* profile type. This distinction *exactly* parallels the segregation of 4-codon box and 2-codon box designations for serine in typical depictions of the code’s sixty-four codons. Further, 4-codon box anticodon types are distinguishable by: (i) relative percentage gaps between preferred lengths per anticodon triplet differ among the three; (ii) the sum of percentages for each anticodon triplet indicates the relative amounts of less frequently observed lengths (see Tables S3−S4), implying the overall patterns are unique. To confirm (ii), the fourth most frequent lengths are: tRNA^Ser(CGA)^ (86 nt); tRNA^Ser(GGA)^ (87 nt); tRNA^Ser(UGA)^ (82 nt).

There is another perspective from which one concludes tRNA are unique: some anticodons are more prone than others to encode multiple copies within a single genome. When Archaea possess multiple copies of a single tRNA, it is usually twofold, or occasionally threefold. Four or more copies are found only for tRNA^Ala(UGC)^, tRNA^Cys(GCA)^, tRNA^Thr(UGU)^. The full distribution of multiple copies as a function of tRNA anticodon triplet and copy number, with a concise description of the relationship among copies is provided in Supplemental Table 5. Permutations vary slightly with copy number:

- If copy number is 2, three options exist: (i) different lengths; (ii) same length, different sequence; (iii) identical length and sequence.
- If copy number is 3, six options appear among these Archaea: (i) different lengths; (ii) same length, but different sequences; (iii) same length and identical sequences; (iv) same length, with two containing the same sequence, while the third copy differs in one or more nucleotide positions; (v) two lengths plus all different sequences; (vi) two lengths, with those that are the same also identical in sequence, while the third differs due to length variance.
- If copy number is 4, eight options are seen: (i) all same length, yet all sequences differ; (ii) all same length, and sequences are exact replicates (clones) of each other; (iii) all same length, plus three copies sequentially identical while the fourth is coded differently; (iv) same length, with two sequences exact duplicates, while the other two are sequenced differently not only from the pair, but from each other as well; (v) two lengths, but all different sequences; (vi) two lengths, with three versions in possession of the same length plus identical sequences, while the fourth copy has a different length; (vii) two lengths, with two versions possessing the same length plus identical sequences, while the other two sequence differently both from the pair and each other; (viii) three tRNA lengths, with two versions possessing the same length and identical sequences, while the other two exhibit different lengths both from each other and the sequence-identical pair. There is no example of a tRNA molecule being encoded in four varieties with all four presenting different lengths.

Copy numbers of five, six, eight exist for tRNA^Cys(GCA)^. Mathematically, more copies demand a greater number of theoretical alternatives with respect to same vs. different relationships. As a practical matter among these Archaea, the five copy cases (five occurrences) appear in three variations; six copy cases (two occurrences) in two versions; the single eight copy case obviously generates one format (see Table S5).

The number of times multiple copies are encoded goes from zero for tRNA^Arg(CCU)^, tRNA^His(GUG)^, tRNA^2Ile(C+AU)^, tRNA^Pro(CGG)^, tRNA^Pro(GGG)^ to forty-six for tRNA^Ala(UGC)^. An exhaustive list of tRNA with twenty or more multiple copy events, besides tRNA^Ala(UGC)^, includes: tRNA^Cys(GCA)^ (41), tRNA^Glu(UUC)^ (37), tRNA^Asp(GUC)^ (33), tRNA^iMet(CAU)^ (33), tRNA^Gly(GCC)^ (31), tRNA^Leu(GAG)^ (30), tRNA^Asn(GUU)^ (27), tRNA^Phe(GAA)^ (27), tRNA^Val(GAC)^ (26), tRNA^Ile(GAU)^ (24). All remaining tRNA anticodon triplets encode 1−19 multiple copy events across these 186 archaeal organisms. Triplets most prone to yielding multiple copies cover the full spectrum of heretofore presumed redundant amino acid codons:

- singleton Met
- doubly degenerate Asp, Asn, Cys, Glu, Phe
- thrice degenerate Ile
- fourfold degenerate Ala, Gly, Val
- sixfold degenerate Leu

It may or may not be biochemically significant that acidic and hydrophobic residues are present among these eleven, but basic residues are absent. Instead, they uniformly constitute the zero frequency category: arginine, histidine, proline. Only two lysine anticodons escape having a representative in this subdivision.

Of greater importance for the principal argument that no code degeneracy exists is the percentage of multiple copies depicting different tRNA lengths or sequences altogether. Either situation challenges presumed redundancy. Since occasions with four or more copies apply to just three anticodon triplets and lead to diverse outcomes with potential ambiguity in interpreting between-copy relationships, conditions with 2−3 copies only per anticodon triplet is considered for this analysis. Eleven anticodons presenting the largest number of multiple copy events (355, 69.5%) are most pertinent for the main thesis since they have more influence on the overall trend. Approximate percentages of different lengths or sequences, as opposed to copies identical in both, are: tRNA^Ala(UGC)^ (2%), tRNA^Cys(GCA)^ (81%), tRNA^Glu(UUC)^ (11%), tRNA^Asp(GUC)^ (15%), tRNA^iMet(CAU)^ (9%), tRNA^Gly(GCC)^ (47%), tRNA^Leu(GAG)^ (63%), tRNA^Asn(GUU)^ (22%), tRNA^Phe(GAA)^ (11%), tRNA^Val(GAC)^ (19%), tRNA^Ile(GAU)^ (63%).

In total, copies differ somehow 27% of the time, excluding seventeen ambiguous cases involving three copies plus nineteen occasions for 4−8 copies uncounted among the eleven. Superficially, this seems weak support for a nonredundancy hypothesis. However, a substantial number (frequency) of nonidentical copies for an anticodon triplet leads to a lack of uniformity *internal* to that molecular type. Since these eleven correlate to eleven amino acids, and tRNA outside this set refer to the same amino acids as these eleven for alanine, glutamic acid, glycine, isoleucine, leucine, valine, one concludes a lack of *external* uniformity across anticodon triplets exists as well.

Nonidentical copies in two formats (length or sequence) may both yield biochemically relevant information. The missing 30.5% yield higher percentages of nonequivalent copies overall, because copy differences in all 511 events equals 44% for two-copy versions and 46% for three-copy occasions, not including seventeen ambiguous cases and twenty occurrences where copy number exceeds three. Hence, a proposition of nonredundancy is even stronger if all available data are considered.

The critical concept is *biochemical information of relevance*. To explore this aspect, removal of identical sequences across archaeal genomes must be undertaken so the residue constitutes every unique sequence for each amino acid-charging anticodon triplet combination. The simplest way to conduct this parsing is to sort all tRNA sequences hierarchically: (i) by amino acid correlate; (ii) by anticodon triplet representing each amino acid; (iii) by length per anticodon. Pairwise comparison of sequences was then performed visually, and not by any available algorithms.

Unique sequences are defined absolutely: single base mismatches at any position between strings is sufficient to declare sequences unique; in other words, only 100% nucleotide-for-nucleotide matches epitomize identicality. For these Archaea, 8869 initial sequences led to 4658 (52.5%) absolutely unique. Supplemental Tables 6−26 (twenty for principal amino acids plus one for selenocysteine, suppressor, and anticodon-indeterminate sequences) provide specifics; amino acids having multiple anticodons arranged under separate tabs within a single Table.

Determining unique sequences per anticodon places an upper bound on a possible set of distinct messages. Tables S6−S26 allow the practitioners of biochemical and microbiological arts to expeditiously find whatever codes-within-the-code potentially exist. Hidden codes should convey the full information content and contextual meaning for each tRNA nucleotide string. Tables S6−S26 also allow assessment of distribution by length profiles per anticodon triplet. Supplemental Table 27 directly descends from Table S3: the latter applies to 8869 sequences, while Table S27 restricts that information to 4658 unique types. Table S27 highlights three other pieces of information: (i) percentage of unique sequences relative to total sequences per anticodon; (ii) number of times a nucleotide triplet appears at the 5’ end; (iii) number of times a nucleotide triplet appears at the 3’ end. Table S27 is complex, so it is transformed into Table 4 (item i) and Table 5 (items ii, iii), leaving length distribution data to Table S27. In different ways, they all demonstrate nondegeneracy. From 148 tRNA employing tRNA^Arg(CCG)^ to 290 for tRNA^Cys(GCA)^:

**Table 4.**
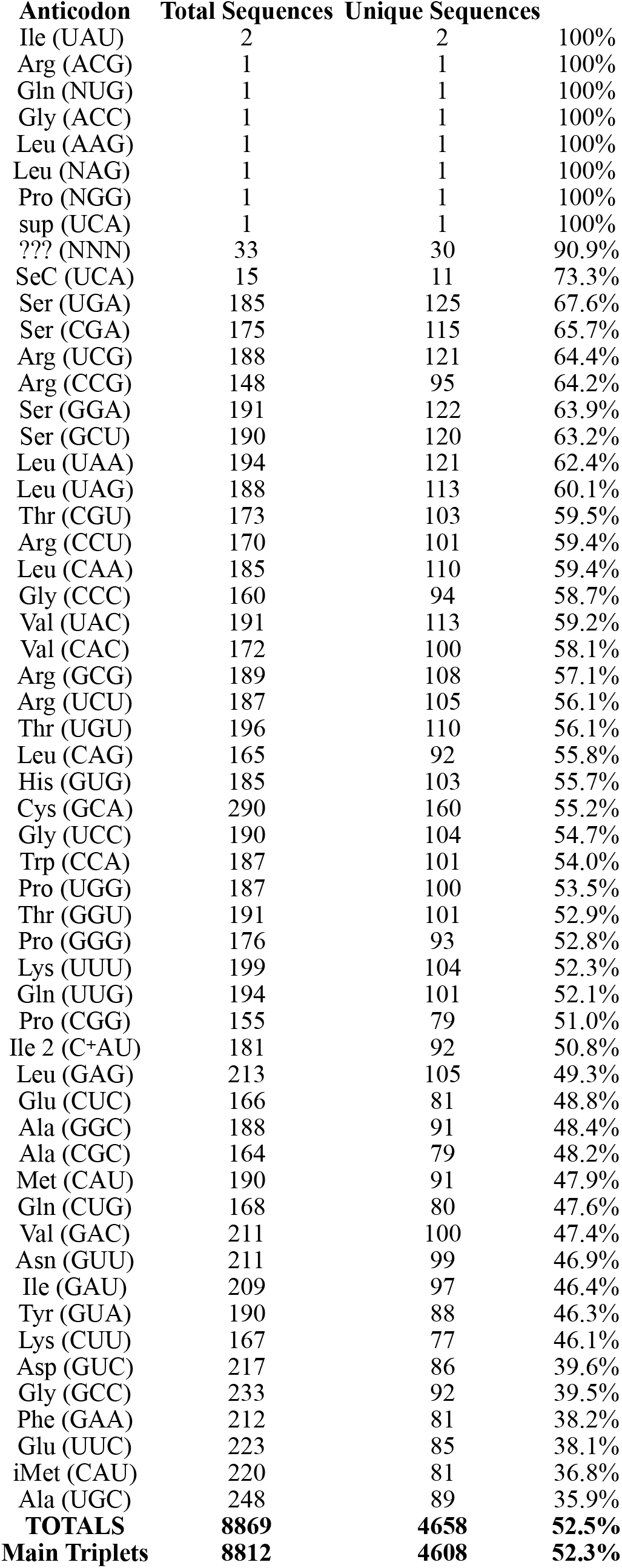
tRNA Unique Sequences per Anticodon.

**Table 5.** First and Last Nucleotide Triplets in tRNA Unique Sequences.

|  |  |  |  |
| --- | --- | --- | --- |
| **First 3 Nucleotides** | GGG 1625 34.9% | **Last 3 Nucleotides** | CCA 2031 43.6% |
| Top 10 = 95.2% | GCC 1241 26.6% | Top 8 = 96.3% | GCA 876 18.8% |
| 33 unique triplets | GCG 727 15.6% | 31 unique triplets | GCG 457 9.8% |
|  | AGC 190 4.1% |  | CCG 453 9.7% |
|  | GCU 189 4.1% |  | GCU 352 7.6% |
|  | GGA 135 2.9% |  | CUA 203 4.4% |
|  | GCA 119 2.6% |  | GCC 62 1.3% |
|  | GUC 79 1.7% |  | GGA 51 1.1% |
|  | CCC 67 1.4% |  | CGA 38 |
|  | AGU 63 1.3% |  | ACG 32 |
|  | GGU 41 |  | UCG 27 |
|  | GGC 40 |  | GUA 24 |
|  | GUU 18 |  | CCU 19 |
|  | AGG 17 |  | GAC 15 |
|  | ACA 15 |  | ACC 10 |
|  | GAC 14 |  | CUG 9 |
|  | UCC 14 |  | ACU 8 |
|  | CCG 12 |  | UCA 5 |
|  | CGG 11 |  | ACA 2 |
|  | GUG 8 |  | CAA 2 |
|  | ACC 7 |  | GGU 2 |
|  | GAG 6 |  | AAA 1 |
|  | CUC 3 |  | AGA 1 |
|  | GUA 3 |  | AUG 1 |
|  | ACG 2 |  | AUU 1 |
|  | ACU 2 |  | CAC 1 |
|  | CGC 2 |  | CAU 1 |
|  | GAA 2 |  | CCC 1 |
|  | UGG 2 |  | CGU 1 |
|  | AGA 1 |  | GGC 1 |
|  | AUC 1 |  | GUU 1 |
|  | GAU 1 |  |  |
|  | UGC 1 |  |  |

- every anticodon triplet expresses a different percentage of unique sequences (Tables 4, S27)
- distinct patterns of length exist (Table S27)
- extreme diversity in both 5’ and 3’ termini are present (Tables 5, S27)

To substantiate this last point, 5’ triplets exhibit slightly greater variety than 3’ triplets. There are thirty-three leading triplets: GGG most common (34.9%), GCC next (26.6%), GCG third (15.6%). The top ten most common account for 95.2% of all variants. By contrast, the final nucleotide triplet is CCA 43.6% of the time, and the most prevalent ending string; GCA is relatively frequent (18.8%), as are GCG (9.8%) and CCG (9.7%). Overall, thirty-one triplets are found, with the top eight accounting for 96.3% of unique tRNA sequences.

Table 4 reports, but excludes from subsequent analysis, possible patterns in: (i) anticodons from sequences (by definition unique) in 1−2 organisms, such as tRNA^Ile(UAU)^; (ii) three cases of incomplete sequences from *Thermococcus A litoralis* necessarily treated as novel; (iii) sequences not attributed to specific anticodons in GtRNAdb; (iv) selenocysteine tRNA^Sec(UCA)^. Omission produces 8812 sequences from the original 8869 involving forty-six anticodons linking twenty amino acids, and unique sequences constitute 4608 (52.3%). Said differently, 47.7% of all encoded sequences are *nucleotide-for-nucleotide identical* throughout their entire length, regardless of that length, with one of the designated unique ones. These figures are reasonably consistent with the multiple copy data, whereby 44% (two copies) and 46% (three copies) depict nonidentical sequences over all tRNA lengths.

Frequent depiction of tRNA in their 2D cloverleaf arrangement makes it more difficult to uncover pertinent observations. If tRNA sequences are arranged linearly left to right (i.e., 5’ → 3’), the amino acid attachment point is on the right. A centerline drawn vertically equals the approximate midpoint for each extended sequence. For bases A/C/G/U written in *Times New Roman* font, the centerline is merely an estimated midpoint because characters are not spaced equally. When same-length sequences are stacked, anticodon triplets indicating the translated amino acid frequently fail to possess perfect vertical symmetric alignment: inserted nucleotides to the left of the triplet cause *rightward skew*; inserted nucleotides to the right induce *leftward skew* (see Tables S6−S26). Based on vertical alignment, it is proposed that the range for tRNA sequences amenable to translation is 70−92 nt, which means the centerline is approximately at positions 35−46 proceeding from the extreme left. This suggestion is slightly discordant with 74−95 nt

(Heinemann *et al*. 2010) and 70−100 nt (Fujishima and Kanai 2014) found among other estimates in the literature. With one exception, all anticodon triplets appear left of, or contiguous to, that centerline: extending to the right is tRNA^Arg(CCG)^ (79 nt). For sequence lengths above or below 70−92 nt, anticodons may appear anywhere along that horizontal line.

Some anticodon triplets are ambiguous because adjacent nucleotides extend the triplet domain. A complete set of multi-position possible anticodon triplets over all encoded tRNA lengths consists of:

- tRNA^Gly(CGC)^ appears as CGCGCGC six times out of seventy-nine unique sequences
- tRNA^Glu(CUC)^ appears as CUCUC in 98% of unique sequences
- tRNA^Gly(ACC)^ is ACCACC in its only sequence
- tRNA^Gly(CCC)^ is CCCCCC instead of CCC in 11% of unique sequences
- tRNA^Lys(UUU)^ appears as UUUU with 100% frequency
- tRNA^Pro(GGG)^ appears as 4−7 consecutive G (i.e., GGGG → GGGGGGG) with no simple triplet
- tRNA^Val(CAC)^ is CAC (20%), CACAC (77%), or CACACAC (3%)

As Table 4 reveals, ignoring anticodon triplet tRNA from a single organism (100%), unassigned sequences (91%), and tRNA sequences associated with selenocysteine (73%), eight anticodons possess sequence uniqueness percentages above 60%, with each referring to an allegedly sixfold redundant amino acid: tRNA^Ser(UGA)^ (67.6%), tRNA^Ser(CGA)^ (65.7%), tRNA^Arg(UCG)^ (64.4%), tRNA^Arg(CCG)^ (64.2%), tRNA^Ser(GGA)^ (63.9%), tRNA^Ser(GCU)^ (63.2%), tRNA^Leu(UAA)^ (62.4%), tRNA^Leu(UAG)^ (60.1%). The opposite extreme yields six triplets with under 40% unique-to-total sequence ratios. They are sufficiently diverse that no pattern with respect to amino acid properties is discernible: tRNA^Ala(UGC)^ (35.9%), tRNA^iMet(CAU)^ (36.8%), tRNA^Glu(UUC)^ (38.1%), tRNA^Phe(GAA)^ (38.2%), tRNA^Gly(GCC)^ (39.5%), tRNA^Asp(GUC)^ (39.6%).

One purpose for establishing a compendium of unique sequences is to confirm or reject the view that for many amino acids, the codons represented are genetically and biochemically redundant. It is now possible to challenge that dogma and place it on trial. The question is: how many sequences of supposedly interchangeable anticodons referring to one amino acid are identical? The answer is *very few*. Evidence against the proposition of degeneracy is the purpose of Supplemental Table 28, but prior to explaining its contents, features concerning all three domains as well as some Archaea-related clarifications are needed.

First, the genetic code for Archaea, Bacteria, Eukarya cites tryptophan as always translated by an inherently nondegenerate codon, meaning no comparisons between anticodons are possible. Second, all three domains recognize methionine initiating protein sequences demands responsibilities fundamentally different than methionine elongating protein sequences. This distinction is highlighted in absolute terms in Archaea: all iMet tRNA begin with adenosine and all eMet tRNA begin with guanosine. Third, other amino acids correlate with single archaeal tRNA besides tryptophan, also rendering anticodon comparison impossible. This applies to asparagine, aspartic acid, cysteine, histidine, phenylalanine, tyrosine.

Alanine, arginine, glutamine, glutamic acid, glycine, isoleucine, leucine, lysine, proline, serine, threonine, valine possess multiple anticodons in Archaea permitting sequence comparisons across unique nucleotide strings. Table S28 is unambiguous and incontrovertible: *no identical sequences* appear for Ala (3 triplets), Gly (3 triplets), Ile (3 triplets), Pro (3 triplets), Ser (4 triplets), Thr (3 triplets), Val (3 triplets). On the other hand:

- Arg (5 triplets) divides into Arg^(CCU)^ and Arg^(UCU)^ plus a grouping for Arg^(CCG)^, Arg^(GCG)^, Arg^(UCG)^ as expected for two-codon box plus four-codon box arrangements in the code. For the set of two anticodon triplets, there exists one pair (74 nt) and five pairs (75 nt) of identical sequences. No matches result for the three anticodon triplet group, therefore none for all five Arg anticodon triplets collectively.
- Gln (2 triplets) yields seven sequences at 73 nt and four at 76 nt with exact matches other than the anticodon triplets.
- Glu (2 triplets) offers one pair of identical sequences at 73 nt, five pairs at 75 nt, and two pairs at 78 nt outside the anticodon triplets.
- Leu (5 triplets) also divides into two-codon box and four-codon box subsets. In contrast to Arg, no identical sequences appear for Leu(^CAA)^ and Leu^(UAA)^, but one set at 85 nt plus three sets at 88 nt exist for Leu^(CAG)^, Leu^(GAG)^, Leu^(UAG)^. Consequently, no sequence matches are discovered applicable to all five anticodon triplets.
- Lys (2 triplets) presents one pair of identical sequences at 74 nt and another pair at 77 nt, excluding the anticodon triplets.

Thirty-one cases involving sixty-six sequences among five amino acids: twenty-seven two-codon boxes and four four-codon box anticodons align 100% nucleotide-by-nucleotide. Perfect matches derive from 4608 unique sequences (1.4%). Yet, 32.4%−64.1% identity exists *within* tRNA anticodons over the canonical twenty amino acids. It seems impossible to maintain allegiance *prima facie* to any concept of codon degeneracy in the genetic code.

## DISCUSSION

It is certainly possible to assert: (i) the expansive scope of tRNA lengths are without biochemical import; (ii) all tRNA for a given amino acid necessarily express degeneracy simply *because* they charge the same amino acid; (iii) multiple lengths per anticodon triplet are benign examples of random insertions or deletions. However, one is then forced to prove all variations in nucleotide number are of sufficiently recent origin such that organisms have had insufficient time to extract them from the reservoir of tRNA; or they represent examples of cost/benefit calculations, whereby each archaeal species, without exception, independently concludes excisions are not worth the energetic costs. It is improbable either argument can be rationally sustained in the face of dramatic divergence among supposedly degenerate anticodons.

It is difficult to justify contrasting positions *vis à vis* tRNA sequence identity. If anticipating high fidelity as normal given the sensitive nature of a translation process for which severe consequences follow from subsequent unintended error, then extensive sequence divergence among them for a single anticodon triplet must be deemed significant, with potential biochemical repercussions needing to be deciphered. If tolerance for high degrees of sequence variance is regarded as standard when accounting for many species and their diversity of habitats, then 32.4%−64.1% exact identity must be rationally explicated in light of expected evolutionary divergence over time. To accord exactitude to high frequencies of horizontal gene transfer by common progenitors as explanation for identity seems an implausible hypothesis demanding confirmation.

The range of tRNA sequences considered here does not include chemically modified nucleotides beyond C^+^ for tRNA^2Ile^. If they were included, then they would provide additional options beyond those cataloged in Tables S6−S26. This need not imply every nucleotide difference between tRNA molecules sharing an anticodon is guaranteed to embody profound biochemical importance, though it is possible this is the case. The single mRNA base change leading to presence or absence of sickle cell phenomenon is a possible precedent (Clancy 2008). All that can be logically stated is single nucleotide variance between two tRNA with the same anticodon cannot be *assumed* irrelevant.

It is conceivable chunks of consecutive base alteration translationally express divergent meaning. This idea is deemed plausible when it is recognized that every tRNA sequence contains bursts of same nucleotide repetition from 2−7 times involving all four bases. Disrupting patterns such as AAA, CCCCC, GGGGGGG, or UUUU may have major impact on translation. It is even possible a tRNA chain sequence taken as a whole, independent of length, allocates real meaning. Lastly, length itself could convey crucial information to be interpreted, and sequences, except for anticodons, are mere camouflage. All options are on the table to be explored by diligent future investigation.

Some nucleotides must be invariant to restrict and maintain 3D topology (Marck and Grosjean 2002). Their study investigated fifty sequences, but just thirteen for Archaea. Considerably more archaeal tRNA have been sequenced over the last twenty years; it is likely some of their observations regarding structural demands need revision. Furthermore, although the overall need for fixed nucleotides in order to have structural integrity remains, given the extreme variation in tRNA lengths shown here, the nature of which bases are conserved may vary between individual species. If true, no general model might apply.

Historically, the focus in comparative genomics research has been on conserved bases or amino acids. This is the underlying strategy for compilations found in databases such as *Clusters of Orthologous Genes* available from NCBI, as well as the *Kyoto Encyclopedia of Genes and Genomes* (KEGG; https://www.genome.jp). Digital information theory, however, postulates interpretational meanings derive from sequence differences, not their constancy (Shannon and Weaver 1949). This point of view offers inherent justification to suggest what may hold for electronic bits might also be germane to an A/C/G/U tetra-code. Ecological principles operate on the same idea: differences in phenotype equals adaptability by genotype to altered circumstances. It would be ill-considered to ignore potential meanings hidden within complete sequences of nucleotides, especially since disguised information concerning element isotopes is scientific precedent.

What potential biochemical consequences are in play if there truly is a code-within-the-code to be uncovered? Transfer RNA could be partially responsible for: (i) amino acid modification during (or after) protein formation; (ii) induction of secondary and/or tertiary structural constraints for mature proteins. In support of the latter notion, Biro (2012) proposed tRNA is instrumental in protein folding. For either or both options to become feasible, it is theorized tRNA sequences bind additional chemical agents designed to tag the growing protein chain at ribosomal A-sites and/or P-sites. To accomplish this task, timing for detachment and removal of tRNA from ribosomal complexes after amino acid charging would be critical factors in determining whether coded messages are viable conduits for these types of exchanges. Based on codon usage patterns, there is evidence tRNA dissociate from interactions with ribosomes at rates slower than the kinetics of translation (Cannarozzi *et al*. 2010). This could mean they stick around long enough to assist with other molecular reactions, such as amino acid alterations.

Among amino acids undergoing co-or post-translational modification, indisputable is disulfide bridge formation by cysteine. Could instructions for S—S bond formation, to the exclusion of alternate linkages, be revealed by nucleotide sequences in tRNA? Is it coincidence that tRNA^Cys(GCA)^ is encoded most often (290 times) in this collection, and presents the highest percentage of nonidentical multiple copy occasions (81%) among the most multiple copy-prone anticodon triplets? Does this not reasonably suggest a potential wealth of information to be extracted from its sequences?

Table S10 contains 160 unique sequences for tRNA^Cys(GCA)^ from 290 encoded (55.2%); of these 160, the most frequent length is 72 nt (39%). Table 6 Set A shows two 72 nt strings displaying differences in 47% of their positions. Does it not seem plausible that this degree of variance has some repercussions *vis à vis* directions for S—S linkage? A need for experimental verification, or hypothesis rejection, is of crucial importance for understanding the full role of tRNA molecules in translation beyond providing an anticodon triplet and readying amino acids for transport.

**Table 6.**
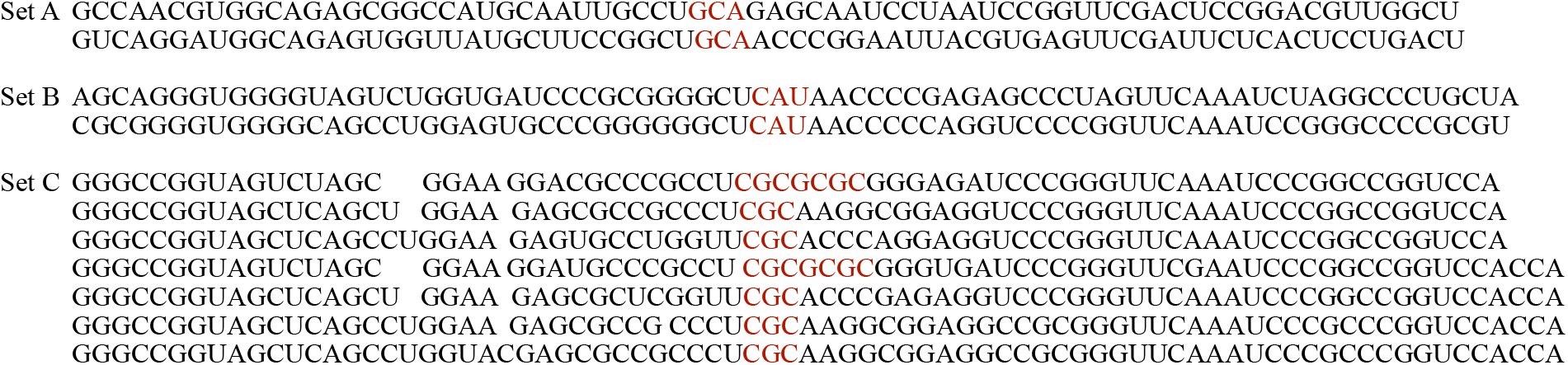
Representative Sequence Comparisons.

*In vivo* active proteins are not formed merely as a consequence of stringing one amino acid after another. A host of secondary and tertiary interactions need development in order for molecules to assume functional end states. Are instructions for structural features (α-helices, β-sheets, intramolecular hydrogen bonds) encoded in tRNA? If present, by what mechanisms are programmed folding elements transmitted? What additional chemical agents must be in, or adjacent to, ribosomal complexes in order to render specific directions successful?

Different possibilities arise if inquiry is made into roles played by tRNA length in contrast to any concentration on sequence *per se*. Could the number of nucleotides refer to thermodynamic properties dependent upon external conditions (temperature, pH, salt cation concentration) or whether species prefer aerobic or anaerobic conditions for survival and growth (Botzma and Margalit 2011)?

The accumulated data acknowledge amino acids have specialized functions within Archaea, and it is these operational qualities which may help to explain corresponding tRNA features. Beyond a potential cysteine impact already entertained, the most well-defined use is for initiator methionine. In keeping with special structural restrictions necessitated by factor binding, tRNA^iMet(CAU)^ is the penultimately conserved collection of unique sequences (36.8%). Yet, composition variations of notable magnitude are available. The version most often found shows a length of 75 nt (53%), and Set B of Table 6 presents tRNA strings with 24% variance. This level of diversity may seem acceptable in some situations and of no significance. However, 63.2% of tRNA^iMet(CAU)^ are identical independent of length (Table S18), hence 24% deviation between two unique versions is huge. Does this difference have biochemical impact during construction of proteins? It is only by experiment that answers can be ascertained.

The degree of unique sequence diversification by tRNA^iMet(CAU)^ is exceeded only by tRNA^Ala(UGC)^ (35.9%), an outcome due to the large number of identical multiple copies across all species. The reason(s) for such an extreme lack of sequence variation is (are) unknown, though they may be tied to two facts: (i) alanine is the simplest chiral amino acid discovered on Earth from meteoritic impacts (Glavin *et al*. 2020), or synthesized by prebiotic methods (Breslow 2011); (ii) tRNA^Ala(UGC)^ is the source introducing alanine into proteins created by mitochondria of phenotypically diverse Eukarya—birds (*Gallus gallus*), fungi (*Saccharomyces cerevisiae*), insects (*Drosophila melanogaster*), mammals (*Homo sapiens*), and reptiles (*Alligator mississippiensis*)—based on a mitochondrial tRNA source (http://mttrna.bioinf.uni-leipzig.de; Jühling *et al*. 2009).

A different alanine tRNA gives some evidence for speculating high sequence similarity between shorter and longer lengths encoded by different species implies responses to similar intracellular aspects, but variable extracellular elements. Using a hand-picked set of tRNA^Ala(CGC)^ to illustrate the point, Table 6 Set C shows how a pattern repeats with 72−78 nucleotides (spaces added to facilitate comparison). This example offers another simple observation: greater amounts of sequence variation transpire at the 5’ end prior to anticodon location compared to the 3’ end supplying an amino acid attachment point. Alterations often occur as single nucleotide changes in any purine/pyrimidine permutation or as nucleotide reversals, for example UC → CU. The general phenomenon, though not the specific elements, was used to claim it was evidence each part derived from different genes that later unified into the modern version of tRNA (Fujishima *et al*. 2008).

Sequences of length 75−78 nt found in the Set C example lead into a discussion of tRNA ending with CCACCA. Wilusz *et al*. (2011) proposed this terminus is “targeted for degradation.” Study of three tRNA molecules, one from each domain of life, was undertaken in controlled reactions with the enzyme responsible for adding CCA to the 3’ terminus. The archaeal component contributing to their research was from *Sulfolobus shibatae* (later renamed *Saccharolobus shibatae*), but is not one whose sequences were investigated in these pages (Table S1). Their conclusions cannot apply to encoded 3’-CCACCA. Aside from Table 6 Set C sequences for tRNA^Ala(CGC)^, this closing sequence is ubiquitous for lengths 75−79 nt with anticodons from: (i) all three Ala; (ii) Ile and modified Ile 2; (iii) all three Pro; (iv) Trp; (v) all three Val. It is also contained in 1−2 sequences from leucine-, lysine-, and phenylalanine-associated tRNA. If Wilusz *et al*. are correct, it must be that sequences naturally so encoded are distinguishable from others with CCACCA ends attached to sequences subsequent to transcription through immediate i*n vivo* activity of ATP(CTP):tRNA nucleotidyltransferase. Possible mechanisms for distinction should be sought because it is illogical to waste material producing a molecule only to see it eventually destroyed.

Table 5 stipulates species having tRNA with CCACCA termini are a subset of organisms using CCA as the predominant expressed form in 43.6% of all unique sequences. A long-standing belief about tRNA may be untrue. Yue *et al*. (1996) claimed the amino acid attachment point requires 3’-terminal CCA for recognition to complete the charging process by amino acids. This proposition has been continuously repeated since alleged in a book chapter (Sprinzl and Cramer 1979), Those authors could not have been articulating that overarching assessment with respect to Archaea, which had only been theorized about at most two years prior, when Woese & Fox (1977) used rRNA gene comparisons to declare their existence alongside bacterial and eukaryl domains. Nor could Deutscher (1982) have been referring to Archaea in his review of tRNA nucleotidyltransferase enzyme activity: taxonomic language proffering three domains was not proposed until 1990 by Woese *et al*. Nevertheless, every paper pertaining even tangentially to 3’-CCA in tRNA asserts its existence as essential for translation, citing as sufficient documentary support, without further elaboration, Sprinzl & Cramer and/or Deutscher.

An investigation of references from that Sprinzl & Cramer contribution revealed every relevant report relied on experimental details from either *Escherichia coli* or *Saccharomyces cerevisiae*. In itself, this proves assumed relevance to Archaea as *ipso facto* proper is scientifically unwarranted. Complete genome sequencing of these two species did not appear in the literature until 1996 (Goffeau *et al*.) for *S. cerevisiae* and 1997 (Blattner *et al*.) for *E. coli*, although some sequences had been explored by chemical (enzyme digestion) or physical (X-ray analysis) methods before that time.

Four research groups published aminoacylation studies on a complete set of common amino acids during the 1970s. The earliest (Tal *et al*. 1972) was also first to assert 3’-terminal CCA as essential. This contribution was followed by papers printed back-to-back (Fraser and Rich 1975; Sprinzl and Cramer 1975). The fourth can be found almost immediately afterwards (Hecht and Chinault 1976). All but the last team worked exclusively on *E. coli*, although with different cell types; the last duo included the fungus in addition to the bacterium in their investigation. Unfortunately for an essentiality argument, *E. coli* may be the worst possible species on which to base the proposition. Why? *It is one of the very few species for which every tRNA with every anticodon associated with every amino acid possesses a 3’-CCA terminus* according to GtRNAdb. This uniformity extends to three molecules (74 nt, 81 nt, 97 nt) whose anticodon triplets could not be assigned. Indeed, *E. coli* universally possessing 3’-CCA was known a decade (Zhu and Deutscher 1987) before the remainder of the genome was sequenced.

If all *Escherichia coli* tRNA always encode CCA nucleotides at the 3’ end, then experimental data showing absence of one or more bases prevents substantive quantities of amino acid acylation is proof of nothing other than transcribed DNA is a natural embodiment of what is evolutionarily necessary *for that organism*. Extension to systems in which the initial condition fails to apply (i.e., not all tRNA are encoded with 3’-CCA), and absent further documentation in those other species, is an improper conjectural leap. This type of transfer of relevance would be analogous to assuming every human has blue eyes and blond hair because it was discovered a family of five fits that category.

If all aminoacylation experiments using *E. coli* are moot since no other results were possible, then what about *S. cerevisiae* tRNA *vis à vis* terminal CCA? Yeast possesses approximately three times the number of tRNA (275) than the bacterium; there are an extraordinary number of copies, most but not all exact duplicates, for almost all anticodons. Encoded as singleton molecules: tRNA^Arg(CCG)^, tRNA^Arg(CCU)^, tRNA^Gln(CUG)^, tRNA^Leu(GAG)^, tRNA^Ser(CGA)^, tRNA^Thr(CGU)^. Overall, 48.4% encode 3’-NCA confined to ten amino acids: sixteen alanine (CCA); twenty-one glycine (GCA, UCA); seven histidine (GCA); fifteen isoleucine (CCA, GCA); twenty-one leucine (CCA, UCA); five methionine (GCA); ten phenylalanine (GCA); fifteen threonine (GCA); six tryptophan (UCA); seventeen valine (ACA, UCA).

According to GtRNAdb, yeast tRNA strings are 71−75 nt except for longer leucine- and serine-related molecules. In *E. coli*, lengths are 74−77 nt except for leucine, serine, and tyrosine (oddly, 85 nt). More precisely, 56% of lengths in the bacterial tRNA are 76−77 nt; in fungal tRNA, 62% are 72−73 nt. If ATP(CTP):tRNA nucleotidyltransferase were to add CCA to 3’ termini in yeast, most lengths in these two species would be nearly identical.

A majority of papers from the 1960s−70s related to tRNA from *Saccharomyces cerevisiae* involve tRNA^Phe^ (3’-GCA) (RajBhandary and Chang 1962; Wong *et al*. 1973; Ladner *et al*. 1975; Sprinzl *et al*. 1977). These particular publications focused on (in order of citation): (i) enzymatic digestion in attempts to unravel tRNA primary sequence; (ii) nuclear magnetic resonance comparison of charged and uncharged versions; (iii) X-ray crystal structure at 2.5Åresolution; (iv) variations in aminoacylation after chemical modifications of 3’-terminal C and A nucleotides, including changes to the ribose ring. An accessible set of yeast tRNA-associated references besides those for tRNA^Phe^ pertain to tRNA^Ala^ (3’-CCA) (Holley *et al*. 1965), tRNA^Ser^ (3’-ACG, UCG) (Zachau *et al*. 1966), tRNA^Tyr^ (3’-AGA) (Kucan *et al*. 1971), tRNA^Val^ (3’-ACA, UCA) (Jilyaeva and Kisselev 1970); they all used chemical or physical techniques to explore structure and reactivity. In summary, most yeast-related studies from this period involved tRNA with 3’-NCA termini.

To resolve the question of 3’-CCA and a supposed essentiality for charging tRNA, it is necessary to describe general reaction procedures. The four groups mentioned differ in experimental details beyond cell types, but common features can be ascertained. Given 3’-CCA for all tRNA in *Escherichia coli* cells renders results from their aminoacylation of no consequence with respect to essentiality, only Hecht & Chinault’s method is of immediate interest. Their aminoacylation procedure included a buffer to control pH, potassium and magnesium salts, a [^3^H] or [^14^C] labeled amino acid plus other unlabeled amino acids, ATP, and both tRNA and aminoacyl-tRNA synthetase corresponding to the labeled amino acid. Notably, the system contained ATP(CTP):tRNA nucleotidyltransferase as part of the synthetase solution.

Omission of CTP meant cytosine could not be attached by nucleotidyltransferase action, but the tRNA could have A as ultimate string ending. The investigation’s objective was to determine amino acid preference for (2’-OH, 3’-H)-ribose *vs*. (2’-H, 3’-OH)-ribose analogs as reaction site. Outcomes were recorded in terms of percent at each location relative to natural (2’-OH, 3’-OH)-ribose tRNA. They do not express absolute percentages for any tRNA charging reaction. However, values other than 0% for either analog implies aminoacylation occurs to some degree, and outcomes other than infinity signify reaction on unmodified tRNA also transpires to some degree. According to their data table, only glutamic acid is listed as an uncertain result for both analogs in yeast, whereas aspartic acid and glutamine both receive this designation for the bacterial analogs. *Uncertain* cannot equate to infinite because aminoacylation certainly results on all *E. coli* tRNA; it most likely means there was some error in data collection. It can therefore be concluded *aminoacylation occurs on each unmodified amino acid-related tRNA in yeast*. Since ten of twenty amino acid tRNA (51.6%) encode non-3’-NCA ends, CCA cannot be essential.

*Essential* is rigorously defined (*Oxford English Dictionary*) as *absolutely necessary* or *impossible to do without*. For example, food is *essential* for the survival of living creatures. Debate endlessly which items constitute food, what organisms are to be included among the living, or the nature of conditions denominated as survival. Once *food, living*, and *survival* become circumscribed terms such that some qualities or features of things are included while others are excluded, a declaration of absolute necessity for things included becomes an incontrovertible element for overall validity in asserting “food is *essential* for the survival of living creatures.”

In biology, if *essential* means something other than *absolutely necessary*, then a statement exactly how it is being used is obligatory, along with a rational justification. Professions of CCA essentiality for translation makes it compulsory that unambiguous evidence demonstrating every protein synthesized in every species within Archaea, Bacteria, Eukarya employs 3’-CCA in every tRNA molecule regardless of anticodon assigned to every amino acid. No exceptions allowed; a single violation suffices to reject the thesis. Supplemental Table 29 indicates frequency of 3’-CCA is inflated by five amino acids in which it is nearly universal regardless of tRNA length: Ala (99.6%), Ile + Ile 2 (95.3%), Pro (100%), Trp (99.0%), Val (99.4%). Unsurprisingly, these same types were mentioned as possessing ubiquitous 3’-CCACCA termini. Other anticodon triplets have CCA as amino acid attachment point just 25.8% of the time.

Theories about the translational process and its evolution are dramatically simplified if it can be claimed, as a generally inviolable rule, that tRNA without naturally encoded CCA termini are forced to accept them by invoking ATP(CTP):tRNA nucleotidyltransferase activity. There is nothing special about the enzyme. Cho *et al*. (2007) altered 2−3 residues, enabling it to be converted from 3’-CCA addition to inclusion of 3’-adducts with UTP and/or GTP. Other mutations initiated 3’-poly(A) or 3’-poly(C) addition with tails averaging twenty nucleotides.

Reformulating the consensus view illustrates the difficulty accepting it without lots of evidence in support, which has not been done. Consider an automobile factory making various makes (amino acids) and models (anticodon triplets) in multiple colors (tRNA lengths). For a select few (Ala, Ile, Ile 2, Pro, Trp, Val), all four wheels (CCA-ending tRNA) are attached to each vehicle in one assembly line operation (transcription). For other makes and models (fifteen amino acids plus corresponding anticodon triplets), a single wheel (CCA as attachment point = 25.8%) is installed. Factory owners (community of scientists) demand these vehicles be moved to another building (ATP(CTP):tRNA nucleotidyltransferases) where the remaining three wheels (addition of CCA) are connected before an automobile is driven off and marketed as fully operational (synthesize proteins through translation).

Does this extended metaphor describe a logical, and biologically efficient, process resulting in an economical way to perform? Why would anyone subscribe to this kind of operating plan, especially since a 3’-CCA ending on all tRNA could be encoded if truly crucial for function?

If sufficient evidence of uniqueness has been offered, one implication is that there is no such thing as *a* model tRNA molecule. If this should become true for Eukarya and Bacteria, and Tables 1−2 hint that it might, then any tRNA molecule is a model primarily of itself. Convictions about translation mechanisms, ribosomal interactions, origin and evolution of the genetic code will all need to be rethought and revised where necessary.

Despite an abundance of evidence in support of tRNA anticodon uniqueness presented here, there may be a compelling reason against supposing sequences provide legitimately significant information for Archaea. Most archaeal species encode just one tRNA sequence per anticodon triplet; cumulatively, there are few multiple copy occurrences. Amino acids using that anticodon would have the same modifications, or invoke structural elaborations in mature proteins in the same fashion. Such rigidity is improbable given the range of sizes and functions for synthesized proteins. Of course, codon usage tendencies, particularly for tRNA with more than one anticodon per amino acid, could circumvent this objection. It also need not apply to translation in Eukarya, which feature many more copies per anticodon triplet, *if* it is shown they are not sequence-exact copies.

There remain a multitude of questions this study has uncovered needing adequate responses. Most tRNA lengths are 70−92 nt (99.4%). Are the thirty-four molecules larger than this interval, designated *mature sequences* by GtRNAdb, fully processed from prior intron-containing precursors (Fujishima and Kanai 2014) and thus functionally active in charging amino acids? If so, which ones? This skeptical query is consistent with observing that anticodon triplets in outsized tRNA appear at sequence positions at odds with standard-sized tRNA. If extra-long sequences cannot engage in translation, why are sterile molecules retained in genomes? Similarly, what about the twenty-two tRNA less than seventy nucleotides? Are they active *in vivo*? If not, why are they found in genomes?

In archaeal alternatives to initiator methionine commencing protein building, tRNA^Leu(CAA)^ aligns with mRNA leucine codon UUG. If 70−92 nt tRNA are appropriate for efficient translation, then based on Table S3, assignment of encoded tRNA^Leu(CAA)^ molecules with length 81 ≤ L ≤ 88 are properly assigned in GtRNAdb. Those with lesser (55−76 nt) and greater (96−218 nt) string length might be better allocated to the *undetermined* category. Reassignment would be compatible with anticodon triplet locations far from the midpoint, except perhaps for the 68 nt sequence. Such a change would conform to the more general question of whether outlier lengths are viable for translation.

Alternatively, if some/many/all atypical tRNA lengths result in charged amino acids available for protein formation, then what does their successful application mean with respect to what is known about requirements for the entire translation process? Specifically, what would it signify about ribosomal space accessibility? The spatial extension of a single nucleotide is 0.3−0.6 nm depending upon whether DNA or RNA, chain length, base composition, cation concentrations, as well as measurement methodology (Chi *et al*. 2013). Can 53−227 nt tRNA all fold into the presumably universal L-shaped 3D geometry of 6 nm × 6 nm × 2 nm (Biro 2012)? Can they also fit successfully into the ribosomal space allotted (Blanchard *et al*. 2004)?

Is ribosomal stalling (Charneski and Hurst 2013; Chevance *et al*. 2014) a feature of time needed to implement messages hiding in tRNA sequences, and independent of goodness of fit within ribosomal complexes? Stalling is associated with misframe reading during translation (D’Onofrio and Abel 2014), and the detailed list of ambiguous anticodons offers reasons to believe their existence facilitate occasions for reading errors. Should messages be embedded in sequences, alignment options between mRNA and tRNA could affect binding molecular agents: (i) aminoacyl-tRNA synthetases; (ii) initiation, elongation, release factors; (iii) ATP for amino acid charging and/or GTP for protein elongation steps; (iv) chemical reagents for possible implementation of instructions for post-translational modification of amino acids and/or protein folding. Shifting anticodon location epitomizes skewness when sequences are written in linear format (Tables S6−S26).

Nonequivalent multiple copies points to an issue whose answer might be philosophical (perhaps metaphysical) rather than biological: what agency decides which sequence to use for a given protein? Decisions at DNA/mRNA/tRNA levels fail because they merely specify codon/anticodon combinations, not the precise version selected. The abundance of research into delineation of factors determinative for codon usage tendencies in singular species or applicable across domains, plus the multitude of opinions discussed in the literature, suggests this question has received no universally acceptable answers as yet. Since it has now been shown that tRNA anticodons are differentiable due to unique sequence and length patterns, perhaps exploration of *anticodon* usage tendencies using these variables will shed light on the issue by working from the other side of the codon/anticodon divide. This suggestion is easier said than done; compilation of data presents a formidable, perhaps insurmountable, logistical challenge.

Codon usage statistics from HiveCuts database (https://hive.biochemistry.gwu.edu/cuts; updated September 2021) are available for 174 of 186 Archaea. They utilize 124,876,735 codon/anticodon pairs to translate 435,278 coding sequences. There are 511 multiple copy events for the set of anticodon triplets, of which almost 50% result in nonidentical copies. The problem is more serious for Eukarya. According to GtRNAdb, *Homo sapiens* encode 416 or 429 (depending on the specimen), or approximately seven per anticodon triplet. Multiples cannot all represent identical copies because Table 1 declares humans encode a variety of tRNA lengths for eleven amino acids: alanine, arginine, asparagine, cysteine, leucine, lysine, phenylalanine, serine, threonine, tyrosine, valine. There are 123,378 protein coding sequences just for our species, whereas 174 Archaea represent 3.5 times that total combined. Is the version selection mechanism biological? Can it be discovered? Now extrapolate to other animals and plants collectively denominated as Eukarya, and the issue of tRNA version selection becomes acute, which forces resolution to become compelling.

Research on codon usage bias (Pal *et al*. 2015) offers widely cited quantitative details concerning the genetic code’s operation. Codon usage is the key component in delineating optimal or rare codons. Optimal, or rare, codons are thematic elements appearing in analyses lead to conclusions about translation kinetics (Kolitz *et al*. 2009), ribosome function (Sabi and Tuller 2014), or how a genetic code formed and evolved (Prat *et al*. 2009). Methods determining optimal codons, for example, use a reference set from highly expressed proteins for elucidation of optimality (Satapathy *et al*. 2016). However, the standard is so vaguely worded that basis sets can incorporate a broad range of proteins, yielding inconsistent results.

To offset problems encountered by incompatible interpretations of an umbrella-term like *highly expressed*, Sharp & Li (1987) attempted to standardize the concept through development of a widely used *Codon Adaptation Index*. Popular approval for their approach has spawned offshoots, such as the *tRNA Adaptation Index* (dos Reis *et al*. 2004), which do not use the set of genes, or organisms, from which the original mathematics were constructed. Alterations perpetuate conflicting results. Emery (2010) and the team of Hershberg & Petrov (2009) separately evaluated optimal codons for the same group of sixty-eight Archaea (including strain), all of which are covered here. They disagreed in their assignments 189 times out of 1224 possible outcomes, a 15.4% discrepancy percentage. Theoretical attributions of meaning to an optimal codons concept become clouded. Perhaps looking at tRNA anticodon usage among these sixty-eight organisms would be insightful, if means were developed to accomplish this task.

If each tRNA is unique in sequence and variable in frequency of usage in proteins down to the specific version of a single anticodon, then what are the implications for notions about evolution of the code itself? Is it plausible that after formation of the earliest codes, subsequent codon/anticodon mappings were introduced almost one-at-a-time, with wobble base connectivity, tRNA nucleotide modification, and changes in tRNA length and/or sequence occurring as new situations were encountered requiring novel instructions for protein building? This concept is concordant with finding that the largest percentages of nonidentical sequences are contributed by arginine, leucine, serine amino acids supposedly displaying sixfold redundancy. More unique sequences equals more varied uses equals more codon assignments. An appropriate analog symbolism might be a constructed image: the human homunculus, in which portions of cerebral cortical area are allocated to body regions based on magnitude of electrical nerve activity. If the idea for code evolution adopting this format has merit, then it is thoroughly dynamic in development, completely antithetical to Crick’s *Frozen Accident* (Crick 1966, 1967).

Arginine and serine have sidechains with hydrogen bond donor/acceptor capability, which would be advantageous for establishing secondary structure within completed proteins. Serine is also probably heavily engaged in post-translational modification when, for example, forming an ether linkage to a sugar to generate glycoproteins. Arginine can perform this same action by virtue of sidechain N-glycosylation. Leucine, alongside isoleucine, tops the hydrophobicity scale (Kyte and Doolittle 1982) with fewer post-translational options; its most crucial role might be involvement in protein sections with transmembrane capability (Liu *et al*. 2002). In summary, a need for diverse sequences for the three amino acids is easily comprehensible.

A rationale for anticodon triplets displaying the smallest fractions of nonidentical sequences is far more difficult to conceive. Inclusion of iMet is probably due to initiation factors demanding stringently reproducible sequences for binding. Anticodon triplets for glycine, alanine, phenylalanine do not lend to a ready explanation for their relative conservatism, although a case for alanine has been offered. Inclusion of aspartic acid and glutamic acid are confounding; they too have donor/acceptor hydrogen bonding roles, and are amenable to sidechain esterification post-translationally leading to many bioactive modifications. It seems more intuitive to believe they demand high fractions of unique tRNA sequences rather than low in Archaea.

If any hidden messages encoded in tRNA could be unraveled, a new realm of possible therapeutic benefit appears. Artificial chemically-designed alterations in sequence and length could result in adjusting how proteins are constructed and function. Supplementing natural proteins with proven desirable traits, and circumventing undesirable toxic proteins, genetic or environmentally caused, might become reality. As DuPont so eloquently advertised decades ago: *better living through chemistry*.

All science is built upon what came before, and it is often assumed, without verification of prior evidence, that published documents provide solid foundations for future construction. Reinventing the wheel serves to challenge the known edifice; in effect to verify, or falsify, past data as validly interpreted.

## MATERIALS and METHODS

Transfer RNA from Archaea utilizing conventional *Genus species* nomenclature were obtained by download from GtRNAdb (http://gtrnadb.ucsc.edu; release 19, June 2021). The database lists 217 entries, but thirty-one were rejected because they provided only strain designations instead of recognized species names; consequently, 186 archaeal tRNA sequences in total were studied. In addition, six Eukaryl and six Bacterial tRNA sequences were provided by the same source. When necessary, organisms were renamed according to the Genome Taxonomy database (http://gtdb.ecogenomic.org; version R207, June 2021). In all cases, only tRNA labeled by the database as *mature sequences* were used, meaning each known intron was removed. Individual sequence lengths per anticodon triplet were given by GtRNAdb and these values were accepted as is with no alterations or checks for accuracy.

Sequences from each domain were separately arranged alphabetically initially by organism genus. Archaea were secondarily sorted within genera alphabetically by species when more than one organism from particular genera were involved; this step was not needed for Eukarya or Bacteria. Mature sequences from all three domains were presorted by GtRNAdb first by reference amino acid, using common three letter descriptors, and second by anticodon triplet per amino acid. The length data given by the database for each sequence string offered a further level of discrimination within every anticodon triplet. Among 186 Archaea, 8869 individual sequences covering all species, tRNA molecules, amino acids, anticodon triplets, and lengths were generated.

These 8869 strings were written horizontally with the 5’ end on the extreme left and 3’ end on the extreme right of each row in a spreadsheet, with columns arranged by organism. On 511 occasions, more than one copy of a sequence, based on anticodon triplet, was encoded. Approximately half of these times, they were exact duplicates; the remaining copies differed by sequence, length, or both. All copies always appeared on separate lines.

From this raw collection (Table S2), archaeal sequences for each amino acid anticodon triplet were combined into distinct batches on individual spreadsheets after length segregations were made. The *sort* command on each spreadsheet arranged any given batch alphabetically (A/C/G/U) starting from the extreme left of each string. In this way, the most sequence divergent string pair could always be found by comparison of top and bottom rows. Visual inspection of successive lines enabled detection of adjacent pairs exhibiting 100% nucleotide-for-nucleotide match across the entire length of the two sequences. All but one of these exact matches were discarded, leaving 4658 unique sequences differing by as little as one nucleotide.

Regardless of whether the focus of analysis was on downloaded sequences, multiple copy events, unique sequences, or beginning (5’) and ending (3’) nucleotide triplets, all that was necessary after each arrangement was simple counting of the number of times some feature was found. In no instance was any sophisticated alignment algorithm utilized; at no time was any statistical package invoked.

## SUPPLEMENTAL MATERIALS

Table_S1._Archaea_in_Study. xlsx.

Table_S2._Archaeal_tRNA_Molecule_Sequences_by_Species.xlsx.

Table_S3._Distribution_of_tRNA_Lengths.xlsx.

Table_S4._Percentage_Distribution_of_tRNA_Lengths.xlsx.

Table_S5._Distribution_of_Multiple_Copies.xlsx.

Table_S6._Unique_Ala_Sequences.xlsx.

Table_S7._Unique_Arg_Sequences.xlsx.

Table_S8._Unique_Asn_Sequences.xlsx.

Table_S9._Unique_Asp_Sequences.xlsx.

Table_S10._Unique_Cys_Sequences.xlsx.

Table_S11._Unique_Gln_Sequences.xlsx.

Table_S12._Unique_Glu_Sequences.xlsx.

Table_S13._Unique_Gly_Sequences.xlsx.

Table_S14._Unique_His_Sequences.xlsx.

Table_S15._Unique_Ile_Sequences.xlsx.

Table_S16._Unique_Leu_Sequences.xlsx.

Table_S17._Unique_Lys_Sequences.xlsx.

Table_S18._Unique_Met_Sequences.xlsx.

Table_S19._Unique_Phe_Sequences.xlsx.

Table_S20._Unique_Pro_Sequences.xlsx.

Table_S21._Unique_Ser_Sequences.xlsx.

Table_S22._Unique_Thr_Sequences.xlsx.

Table_S23._Unique_Trp_Sequences.xlsx.

Table_S24._Unique_Tyr_Sequences.xlsx.

Table_S25._Unique_Val_Sequences.xlsx.

Table_S26._Unique_Sec,_sup,_Undetermined_Sequences.xlsx.

Table_S27._Length_Distribution_of_tRNA_Unique_Sequences.xlsx.

Table_S28._Identical_tRNA_Sequences_by_Amino_Acid.xlsx.

Table_S29._tRNA_Length_Distribution_for_CCA_Terminus.xlsx.

